# Unravelling genomic and functional traits of two biocontrol and plant growth-promoting *Pseudomonas* endophytes

**DOI:** 10.64898/2026.08.28.747936

**Authors:** Gustavo Santoyo, Aurora Flores-Piña, Hugo G. Castelán-Sánchez, Valeria Valenzuela-Ruiz, Sergio de los Santos-Villalobos, Debasis Mitra, Olubukola Oluranti Babalola, Mauricio Schoebitz, Ma del Carmen Orozco-Mosqueda

## Abstract

Plant growth-promoting bacterial endophytes represent a sustainable strategy for enhancing agricultural productivity while reducing reliance on synthetic fertilizers and pesticides. This study focused on the genomic and functional characterization of two endophytic bacterial strains, R11F and R19M, isolated from bean and maize roots, respectively. Comparative analyses based on 16S rRNA gene sequences, average nucleotide identity (ANI), and genome-to-genome distance calculations (GGDC) classified both isolates as *Pseudomonas palleroniana*. Comparative genomic analyses revealed highly conserved genomes containing genes associated with plant colonization, phosphate solubilization, stress adaptation, heavy metal resistance, and hydrocarbon degradation. Genome mining further identified 17 and 18 biosynthetic gene clusters (BGCs) in R11F and R19M, respectively, including non-ribosomal peptide synthetases (NRPS), pyoverdine, NRP-metallophores, RiPP-like compounds, arylpolyenes, β-lactones, terpenes, NAGGN, and hydrogen cyanide. Strain-specific BGCs associated with syringomycin and viscosin biosynthesis were identified in R11F, whereas R19M harbored clusters related to asplenin and kolossin biosynthesis. *In vitro* assays confirmed indole production, phosphate solubilization, and siderophore production, as well as the ability of both strains to grow in nitrogen-free medium. Both strains significantly inhibited the growth of *Fusarium oxysporum*, *Phytophthora cinnamomi*, and *Colletotrichum gloeosporioides*. Furthermore, plant inoculation assays demonstrated host-dependent growth promotion, with R11F showing the most consistent improvements in plant growth parameters in tomato, wheat, and lentil. Overall, the integration of comparative genomics and experimental validation demonstrates that *P. palleroniana* R11F and R19M possess complementary traits associated with plant growth promotion, pathogen suppression, saline stress adaptation, and bioremediation.

## 1. Introduction

The increase in the human population and the growing demand for food have become one of the major challenges that must be addressed in the coming years. Projections from the United Nations Department of Economic and Social Affairs estimate that the global population will reach 8.5 billion by 2030 and 9.7 billion by 2050 [1]. Under this scenario, food demand is expected to exceed current agricultural production. On the other hand, crop production and productivity face environmental challenges such as global warming, salinity stress, soil erosion-induced stress, and phytopathogen attacks. Consequently, the use of agrochemicals, including fertilizers and pesticides, has become widespread [2,3].

It has been estimated that approximately 3.7 million tons of pesticides are used annually worldwide to protect crops [3]. Of these, nearly 50% are not absorbed after application and are released into the environment [2]. Once applied to agricultural systems, agrochemicals are transported into surface and groundwater, dispersing throughout ecosystems and ultimately reaching food products. Although these inputs improve crop productivity, their excessive and prolonged use poses risks not only to human health but also to ecosystem functioning and the services they provide.

Among the strategies proposed to promote more sustainable agriculture, the use of plant growth-promoting bacteria (PGPB) has emerged as a promising approach because of their ability to establish intimate and often persistent associations with plant tissues. These bacteria can enhance plant growth and fitness through multiple, often complementary mechanisms, including the production of phytohormones such as indole-3-acetic acid (IAA), siderophore-mediated iron acquisition, biological nitrogen fixation, phosphate solubilization, and the production of other metabolites that improve nutrient availability and plant development [4,5]. Beyond direct growth promotion, PGPB can contribute to plant resilience by modulating host defense responses, improving tolerance to abiotic stresses such as salinity and drought, and producing antimicrobial compounds that suppress a broad range of phytopathogens. These include important fungal pathogens such as *Fusarium*, *Colletotrichum*, *Botrytis*, and *Rhizoctonia*, as well as oomycete pathogens such as *Phytophthora* and *Pythium*, which are responsible for substantial yield losses in numerous crops [6–8].

Another group of more specialized plant-associated bacteria comprises bacterial endophytes, which are capable of colonizing and surviving within internal plant tissues without causing apparent harm to their hosts. Based on their lifestyle, endophytes can be classified as opportunistic, transient, obligate, or facultative. Opportunistic and transient endophytes enter or colonize plants sporadically, whereas obligate endophytes depend largely on their hosts and facultative endophytes can live both within plants and in other environments [4]. Endophytic bacteria have therefore emerged as promising alternatives or complements to conventional agrochemical inputs because of their ability to mitigate both biotic and abiotic stresses [9]. Several studies have shown that bacterial endophytes can increase plant biomass, enhance tolerance to salinity and other environmental stresses, and contribute to the biological control of important fungal and oomycete pathogens, including species of *Fusarium*, *Colletotrichum*, *Botrytis*, *Rhizoctonia*, and *Phytophthora*, to mention but a few [10–13].

Successful colonization of internal plant tissues requires a diverse repertoire of traits, including chemotaxis, flagella and pili, exopolysaccharides, adhesins, lytic enzymes such as cellulases and pectinases, quorum sensing, antibiosis, and mechanisms to evade or modulate plant defense responses [14,15]. Within this context, several species of the genus *Pseudomonas* have been reported as efficient bacterial endophytes because of their ability to colonize diverse plant hosts, produce bioactive metabolites, exert biocontrol capabilities and promote plant growth [5,16,17]. *Pseudomonas* species exhibit diverse plant beneficial traits, including nutrient mobilization, siderophore and phytohormone production, antimicrobial activity, induction of systemic resistance, and enhancement of tolerance to salinity, tolerance to heavy metals, drought, etc. [18–22].

In this study, we performed a taxonomic and comparative genomic characterization of the bacterial endophytes R11F and R19M, including genome mining to identify traits and biosynthetic gene clusters potentially associated with plant growth promotion, stress adaptation, and pathogen suppression. We further evaluated their plant growth-promoting traits *in vitro*, antagonistic activity against three phytopathogens, and effects on plant growth using three crop models. This integrated approach provides insights into the genomic and functional potential of these bacterial endophytes as candidates for microbial inoculants in sustainable agriculture.

## 2. Materials and Methods

### 2.1 Bacterial strains and culture conditions

The R11F and R19M strains were isolated as endophytic bacteria from the roots of maize and common bean plants, respectively, in Michoacán, Mexico. The isolates were purified and preserved in the strain collection of the Genomic Diversity Laboratory. In addition, the phytopathogenic strains *Fusarium oxysporum*, *Colletotrichum gloeosporioides*, and *Phytophthora cinnamomi* were kindly provided by Dr. Rodolfo López Gómez from the Plant Physiology Laboratory at the Institute for Chemical-Biological Research. (Supplementary Figure 1).

### 2.2 *In vitro* characterization of plant growth-promoting traits

Four *in vitro* plant growth-promoting traits were evaluated: indole production, nitrogen fixation, phosphate solubilization, and siderophore production. For this purpose, the strains were cultured in specific media: nutrient broth supplemented with tryptophan for indole production, followed by detection using Salkowski reagent; nitrogen-free bromothymol blue (NFb) medium to assess the ability to grow under nitrogen-free conditions; Pikovskaya medium for phosphate solubilization; and Chrome Azurol S (CAS) medium for siderophore production.

### 2.3 *In vitro* biocontrol activities

Biocontrol activity was evaluated on nutrient agar/potato dextrose agar (NA/PDA) medium. A mycelial plug of each phytopathogenic strain was placed at the center of Petri dishes containing NA/PDA medium and incubated for 24 h to allow initial fungal growth. Subsequently, the bacterial strains were inoculated at a distance of 2 cm from the fungal mycelium, and the plates were monitored for five days to evaluate fungal growth inhibition.

### 2.4 Plant growth assays under growth chamber conditions

For the plant growth promotion assays conducted under growth chamber conditions, *Solanum lycopersicum* seeds were used. The seeds were surface-disinfected by immersion in 70% (v/v) ethanol for 30 s, followed by immersion in 2.5% (v/v) sodium hypochlorite for 2 min with continuous gentle agitation. The seeds were then thoroughly rinsed six times with sterile distilled water to remove residual sodium hypochlorite and germinated in a peat moss substrate. After germination, the seedlings were transferred to individual pots and inoculated with bacterial suspensions prepared in phosphate buffer at a concentration of 1 × 10⁶ CFU. Two inoculations were performed: the first at the time of transplanting into the substrate and the second 15 days later. Four treatments were established, each consisting of five seedlings: (1) a non-inoculated control (C), irrigated only with water; (2) plants inoculated with strain R11F; (3) plants inoculated with strain R19M; and (4) plants co-inoculated with both strains (R11F–R19M). After 21 days, the plants were harvested, and phytometric parameters were determined.

### 2.5 Greenhouse experiments

Wheat and lentil seeds were surface-disinfected with commercial 2.5% sodium hypochlorite, followed by six rinses with sterile distilled water. The only difference in the disinfection procedure was the exposure time to sodium hypochlorite: wheat seeds were treated for 4 min, whereas lentil seeds were treated for 3 min. After surface disinfection, the seeds were germinated in peat moss substrate, and one-week-old seedlings were transplanted into individual pots.

For the greenhouse assays, seedlings were grown in pots containing peat moss substrate and inoculated every 15 days with 5 mL of a bacterial suspension at 1 × 10⁸ CFU mL⁻¹, with the first inoculation performed at the time of transplanting. The greenhouse experiments were conducted from 5 December 2025 to 15 January 2026. Plants were irrigated weekly and supplemented with Leaf Vitality fertilizer.

The same four treatments established in the growth chamber assays were used: non-inoculated control (C), R11F, R19M, and co-inoculation with R11F and R19M (R11F– R19M). The plant growth parameters evaluated included shoot fresh weight, root fresh weight, root length, shoot length, chlorophyll content, and number of leaves.

### 2.6 Genomic DNA Extraction and Sequencing and Assembly

The R11F and R19M strains were independently cultured on nutrient agar (BD Bioxon) at 30 °C overnight. A single colony of each strain was then selected and grown overnight in nutrient broth (BD Bioxon). The resulting overnight cultures were used for genomic DNA (gDNA) extraction. Genomic DNA was extracted using the SDS/proteinase K method, followed by polysaccharide precipitation in the presence of a high salt concentration, as described by Mahuku (2004). DNA quality and quantity were assessed by agarose gel electrophoresis and spectrophotometric analysis using a NanoDrop spectrophotometer (Thermo Scientific). The DNA samples were sequenced using the Illumina platform, and the resulting reads were assembled using SPAdes v4.0. The assembled contigs were deposited in the NCBI database under accession numbers NZ_JBKORM000000000.1 and NZ_JBKQAR000000000.1.

### 2.7 Genomic assembly and taxonomic identification

A *de novo* genome assembly was performed using SPAdes v4.2.0 in “careful” mode to reduce assembly errors for strains R11F and R19M. The resulting assemblies comprised 65 and 64 contigs, respectively. The assembled contigs were ordered using Mauve Contig Mover v2.4.0, with the genome of *Pseudomonas palleroniana* LMG 23076^T^ (GenBank accession: GCA_900105975.1) as the reference genome. The resulting assemblies showed 99.79% similarity to the reference based on 16S rRNA gene sequences. The assembled genomes were subsequently submitted to the EzBioCloud database to identify their closest phylogenetic relatives, using a species-level threshold of >98.7% 16S rRNA gene sequence similarity.

For species-level taxonomic classification, the genomes were compared with closely related *Pseudomonas* strains identified based on >98.7% 16S rRNA gene sequence similarity. Genome relatedness was assessed using several overall genome relatedness indices (OGRIs), including average nucleotide identity (ANI) calculated with OrthoANI, ANIb using BLAST+ v2.2.29, ANIm using MUMmer v3.0, and *in silico* DNA–DNA hybridisation (dDDH) using the Genome-to-Genome Distance Calculator (GGDC) v3.0 with the BLAST+ method.

Genome assemblies of closely related *Pseudomonas* species were obtained from the TYGS taxonomic identification results and included in the comparative genomic analysis. The dataset comprised *P. palleroniana* R11F (JBKORM01.1), *P. palleroniana* R19M (JBKQAR01.1), *P. palleroniana* LMG 23076^T^, *P. petroselini* MAFF 311094, *P. asgharzadehiana* Phyllo_192, *P. nabeulensis* E10B, *P. pergaminensis* Phyllo_220, *P. aylmerensis* S1E40, *P. tolaasii* NCPPB 2192, *P. costantinii* LMG 22119, *P. extremorientalis* LMG 19695, *P. poae* LMG 21465, *P. marginalis* H14, *P. cremoris* WS 5106^T^, *P. kairouanensis* KC12, *P. azotoformans* LMG 21611, *P. canadensis* 2-92, *P. tritici* D277, *P. gelidaquae* IB20, *P. lurida* LMG 21995, *P. fluorescens* DSM 50090, and *P. edaphica* RD25.

Gene clustering, ortholog assignment, and pangenome analysis were performed using Roary v3.13.0. The core genes shared among all genomes were concatenated to generate a core-genome alignment. A maximum-likelihood phylogenetic tree was subsequently constructed using IQ-TREE, with bootstrap analysis used to assess branch support. The resulting phylogenetic tree was visualised using Phandango.

### 2.8 Genome annotation, functional analysis and genome mining

The Proksee v2.4.0 platform, which incorporates the Rapid Prokaryotic Genome Annotation (Prokka) pipeline, was used to generate circular genomic maps of the strains, displaying coding sequences (CDSs), tRNA and rRNA genes, as well as GC content and GC skew profiles. The genomes were additionally annotated using the RASTtk pipeline v2.0 with default parameters. Potential plant growth-promoting traits were predicted using the PGPT-Pred tool available in PLaBAse v1.02. In addition, the genomes were analyzed using antiSMASH v8.0.2 to identify biosynthetic gene clusters (BGCs) potentially associated with the production of secondary metabolites involved in biocontrol and other plant-associated functions.

### 2.9 Statistical analysis

Differences in plant growth-promoting traits between strains were evaluated in triplicate for each assay. The results were recorded qualitatively, except for indole production, which was quantified using a calibration curve. Differences in indole production between strains were analyzed using an independent-samples t-test. For the antagonism assays, significant differences were assessed by one-way analysis of variance (ANOVA), followed by Duncan’s multiple range test, using three biological replicates (n = 3). Similarly, plant growth parameters were analyzed by one-way ANOVA, and mean comparisons were performed using Duncan’s multiple range test at a significance level of α = 0.05.

## 3. Results

### 3.1 Taxonomic affiliation of R11F and R19M strains

Comparative analyses of overall genome-relatedness indices (OGRIs), including 16S rRNA gene sequence identity, average nucleotide identity (ANI), and *in silico* DNA–DNA hybridization (dDDH) calculated using the Genome-to-Genome Distance Calculator (GGDC), were performed to determine the taxonomic affiliation of strains R11F and R19M relative to closely related *Pseudomonas* species available in the EzBioCloud database, using only type strains (Table 1). Both strains showed the highest genomic similarity to the type strain *P. palleroniana* QT, with 16S rRNA gene sequence identity of 99.79%, ANI values of 98.70% for R11F and 98.77% for R19M, and GGDC values of 89.70% and 89.77%, respectively (Table 1).

**Table 1.**
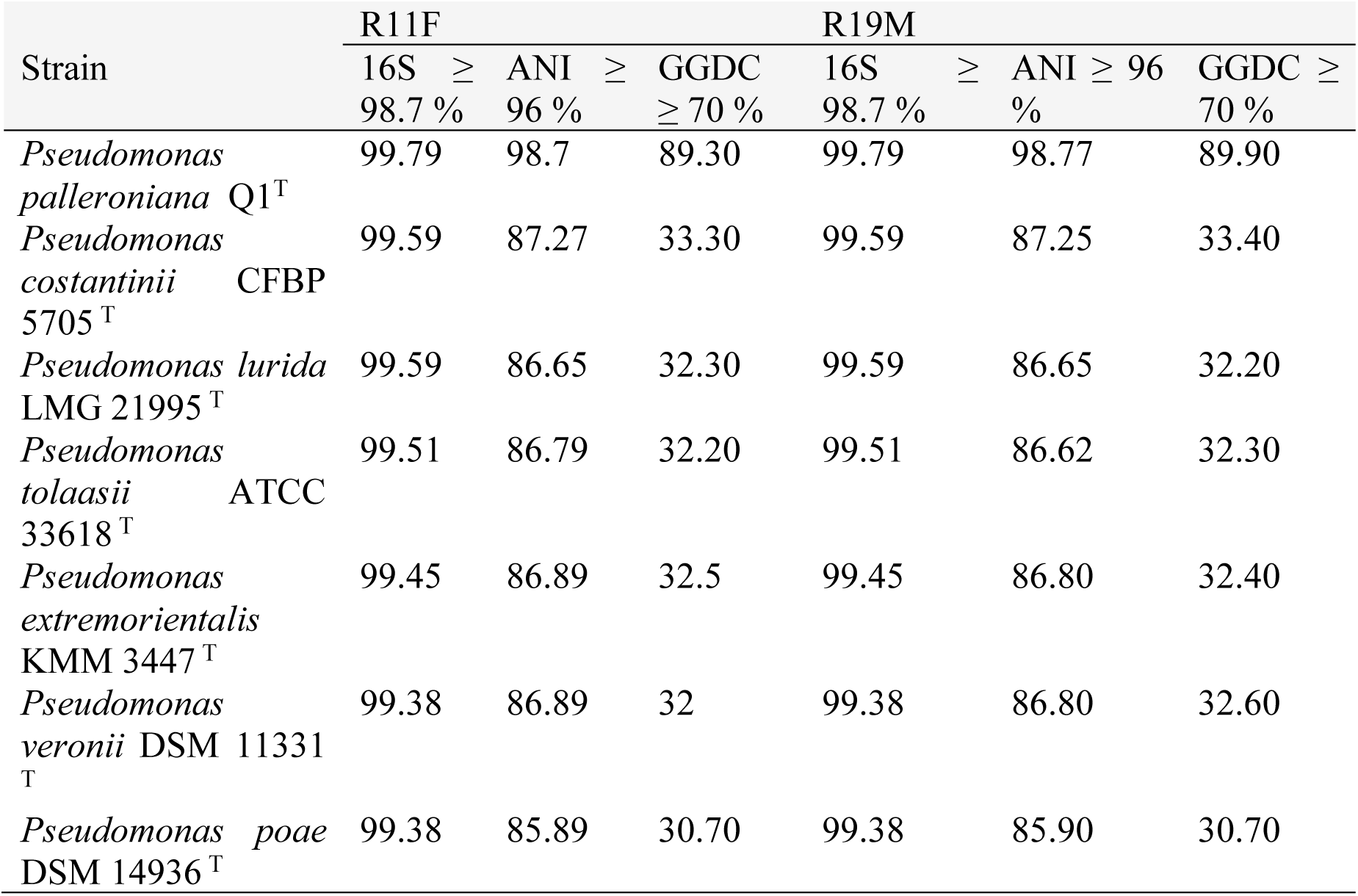
OGRIs of strains R11F and R19M with respect to type species of the genus *Pseudomonas*.

The OGRI values exceeded the established thresholds for bacterial species delineation (ANI ≥ 95–96% and dDDH ≥ 70%), supporting the assignment of both strains to *P. palleroniana*. Although other closely related species, including *P. costantinii*, *P. lurida*, *P. tolaasii*, *P. extremorientalis*, *P. veronii*, and *P. poae*, exhibited high 16S rRNA gene sequence identities, their ANI and GGDC values were substantially lower, indicating clear genomic differentiation from R11F and R19M. These findings highlight the limitations of relying solely on 16S rRNA gene sequences for species-level identification within closely related *Pseudomonas* taxa and demonstrate the value of genome-wide relatedness metrics for resolving their taxonomic relationships.

### 3.2 Phylogenomic relationships, core genome and pangenome analysis

The taxonomic affiliation of the endophytic strains R11F (Figure 1) and R19M (Figure 2) was further confirmed through phylogenomic analyses. Both strains clustered with *P. palleroniana*, forming a well-supported group with the type strain LMG 23076^T^. Given that both endophytic strains were closely affiliated with the same *P. palleroniana* type strain and were isolated from the same ecological niche, we further investigated whether R11F and R19M might represent the same or highly similar genomic lineage. Although both strains were isolated from different plant hosts, they originated from the roots of crops sharing similar ecological conditions. To address this question, we performed core-genome and pangenome analyses of R11F and R19M.

**Figure 1.**
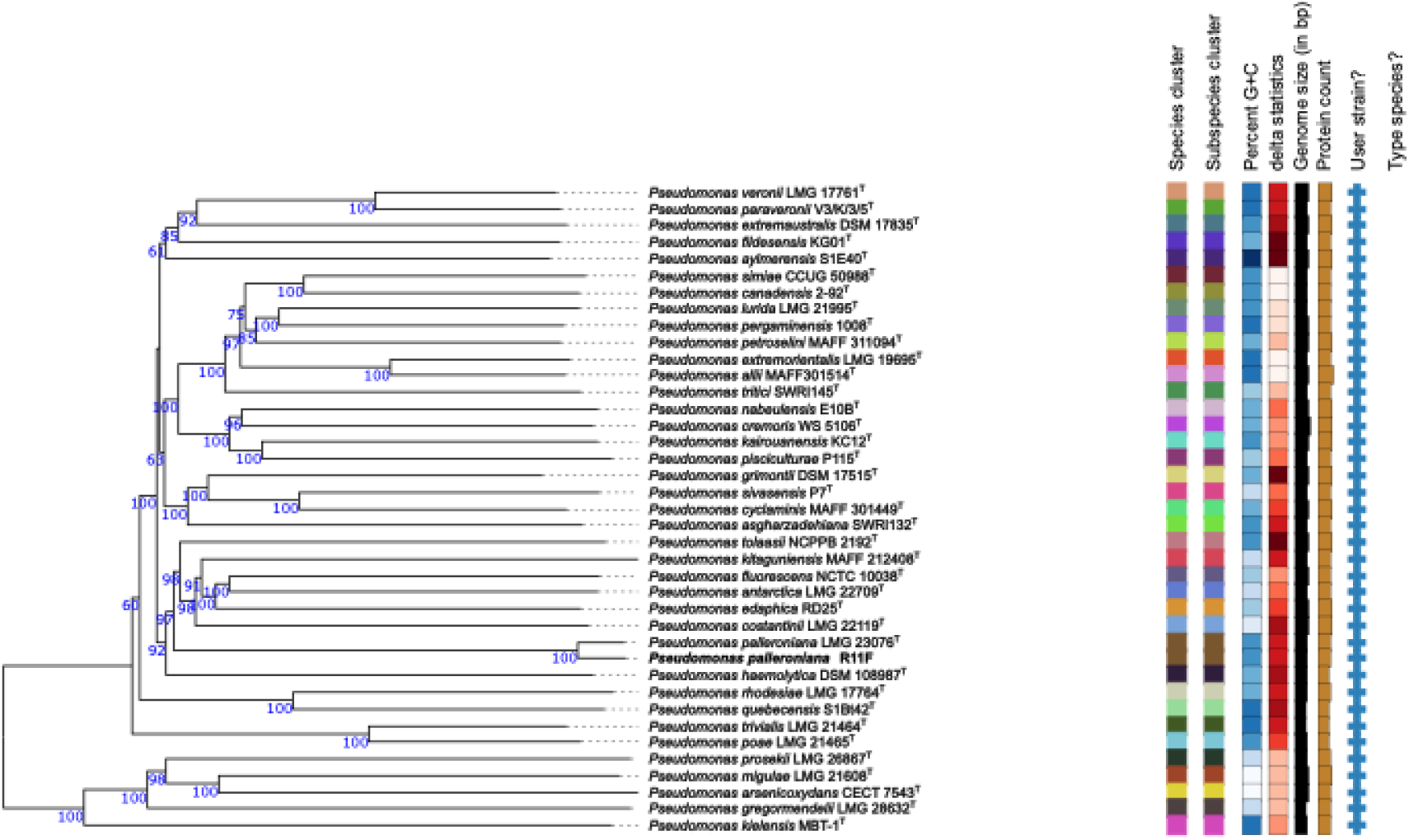
Whole-genome phylogenomic reconstruction showing the position of strain **R11F** relative to closely related taxa in species cluster *a*. The phylogeny was generated through the TYGS platform using FastME v2.1.6.1 and GBDP distances derived from genome sequences (Lefort V, Desper R, Gascuel O. FastME 2.0: a comprehensive, accurate, and fast distance-based phylogeny inference program. Mol Biol Evol. 2015. https://doi.org/10.1093/molbev/msv150.). Branch lengths correspond to d5 GBDP distances, whereas node values represent pseudo-bootstrap support (>60%) from 100 replicates. The average branch support was 89.8%, and the tree was midpoint-rooted (Farris JS. Estimating phylogenetic trees from distance matrices. Am Nat. 1972;106:645–67.).

**Figure 2.**
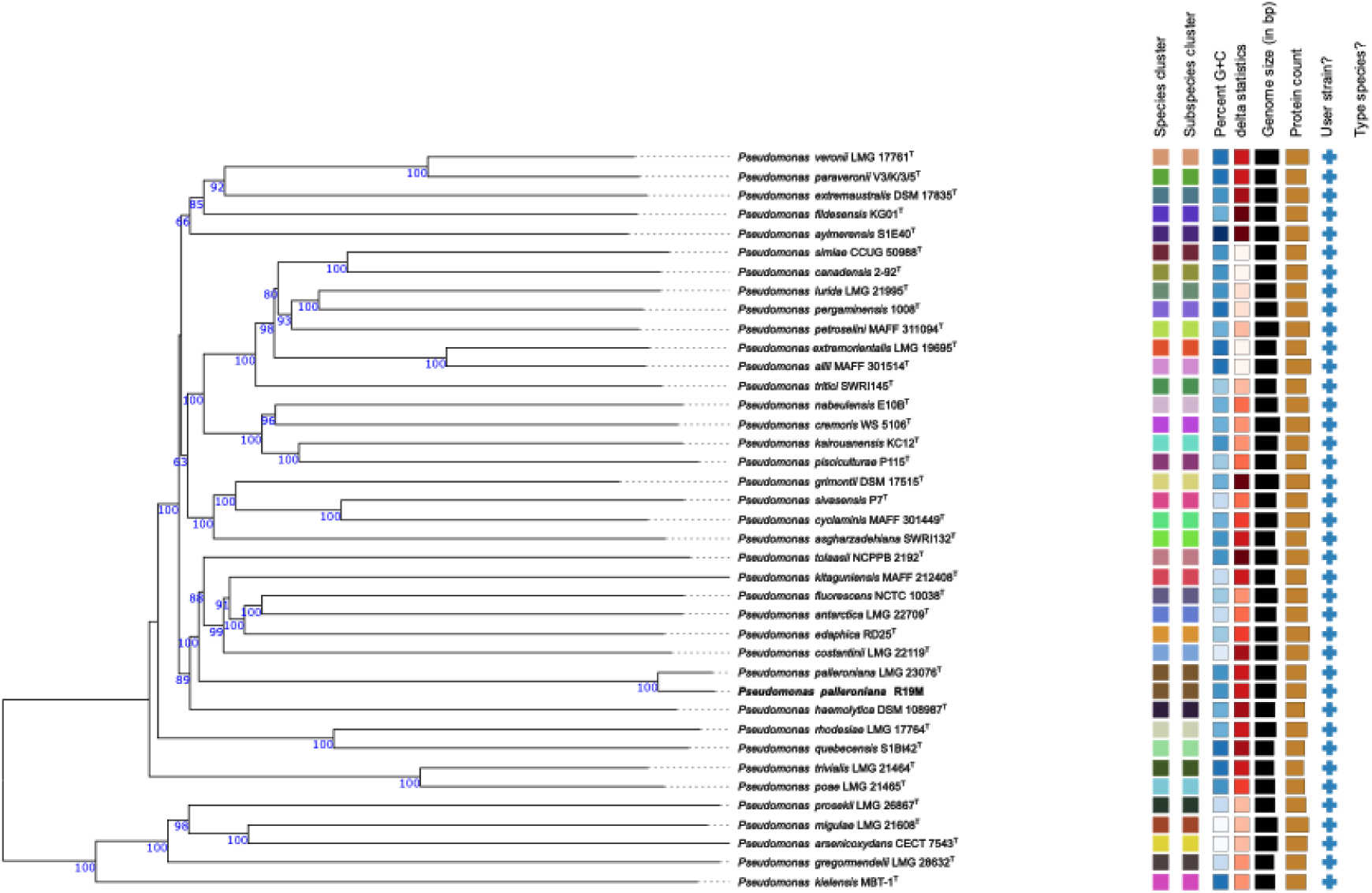
Whole-genome phylogenomic reconstruction showing the position of strain **R19M** relative to closely related taxa in species cluster *a*. The phylogeny was generated through the TYGS platform using FastME v2.1.6.1 and GBDP distances derived from genome sequences (Lefort V, Desper R, Gascuel O. FastME 2.0: a comprehensive, accurate, and fast distance-based phylogeny inference program. Mol Biol Evol. 2015. https://doi.org/10.1093/molbev/msv150.). Branch lengths correspond to d5 GBDP distances, whereas node values represent pseudo-bootstrap support (>60%) from 100 replicates. The average branch support was 89.8%, and the tree was midpoint-rooted (Farris JS. Estimating phylogenetic trees from distance matrices. Am Nat. 1972;106:645–67.).

The results showed that, despite their close phylogenomic relationship, R11F and R19M did not represent the same genomic strain (Figure 3). The gene presence/absence analysis generated using Roary revealed highly conserved core genomes in both *P. palleroniana* strains, with only minor differences in their accessory genes. R11F and R19M exhibited highly similar gene-cluster profiles, consistent with their close phylogenetic relationship. Both strains shared a conserved core genome characteristic of the *P. fluorescens* group while retaining subsets of accessory genes within restricted pangenome intersections (Figure 4). These strain-specific genetic features may contribute to differences in ecological adaptation and plant-associated functions, including plant growth promotion.

**Figure 3.**
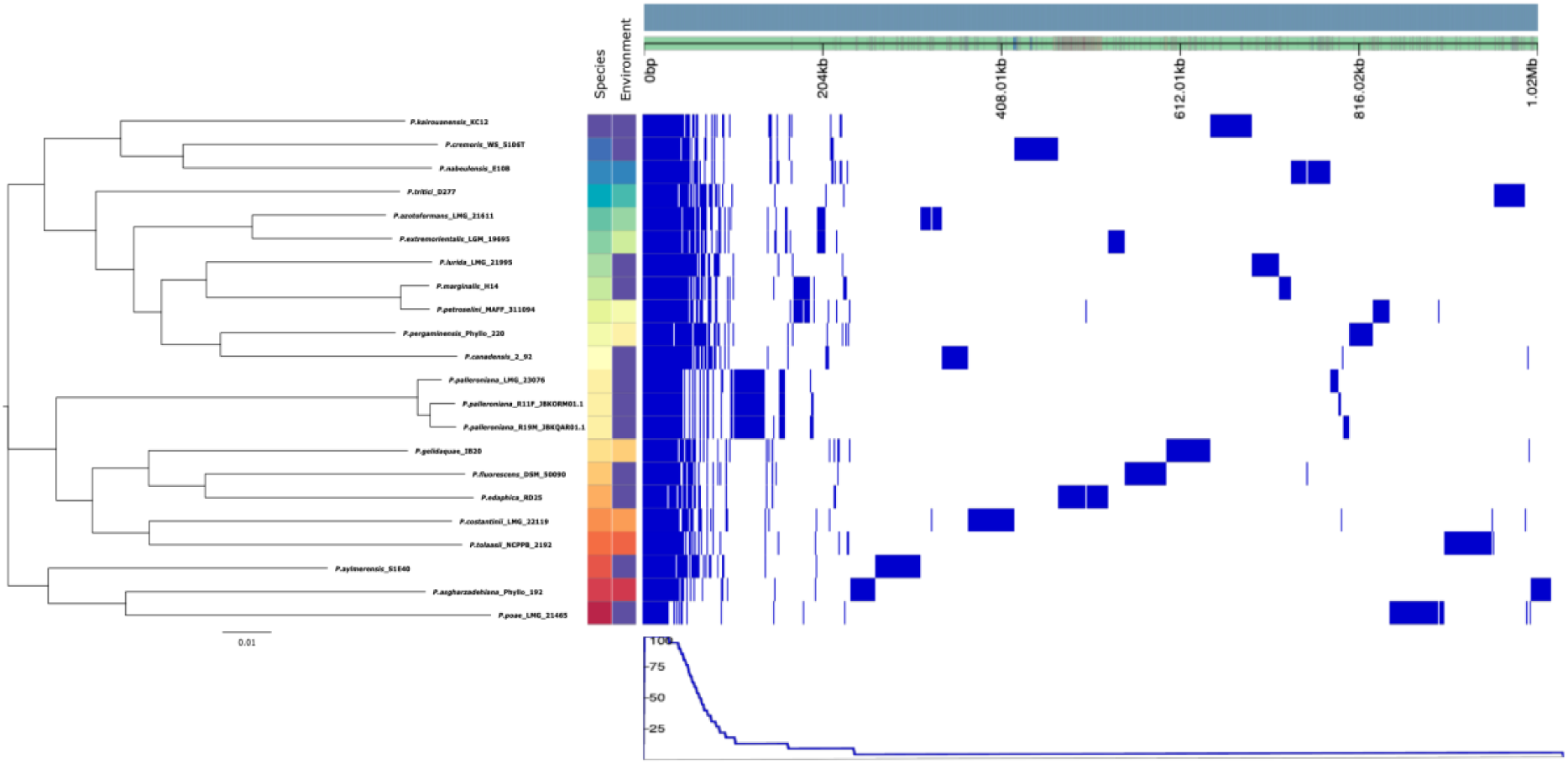
**Core genome analysis of endophytic isolates R11F and R19M among *Pseudomonas palleroniana* and related *Pseudomonas* genomes.**

**Figure 4.**
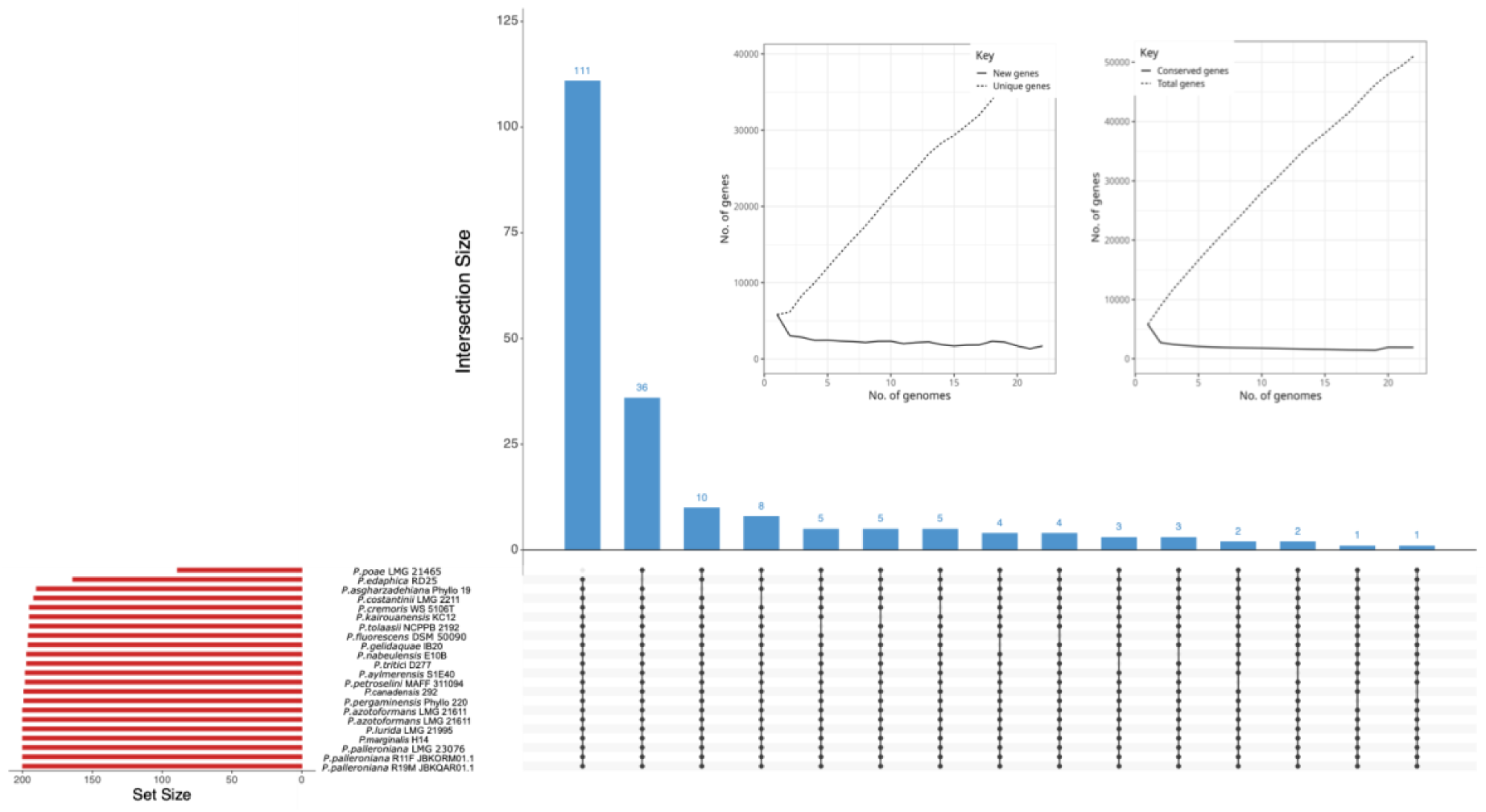
**Pangenome composition and gene conservation among *Pseudomonas palleroniana* and related *Pseudomonas* genomes.**

### 3.3 Genomic traits for plant growth promotion

Functional annotation of the genomes of isolates R11F and R19M using RAST and PLaBAse revealed highly similar functional profiles (Figure 5). According to RAST, the predominant categories were amino acids and derivatives (21%), carbohydrates (11%), protein metabolism (9%), cofactors, vitamins, prosthetic groups, and pigments (9%), fatty acids, lipids, and isoprenoids (5%), and stress response (5%). Both isolates also harbored genes associated with motility and chemotaxis, aromatic compound metabolism, virulence and defense, iron acquisition, and secondary metabolism (Figure 5a,b).

**Figure 5.**
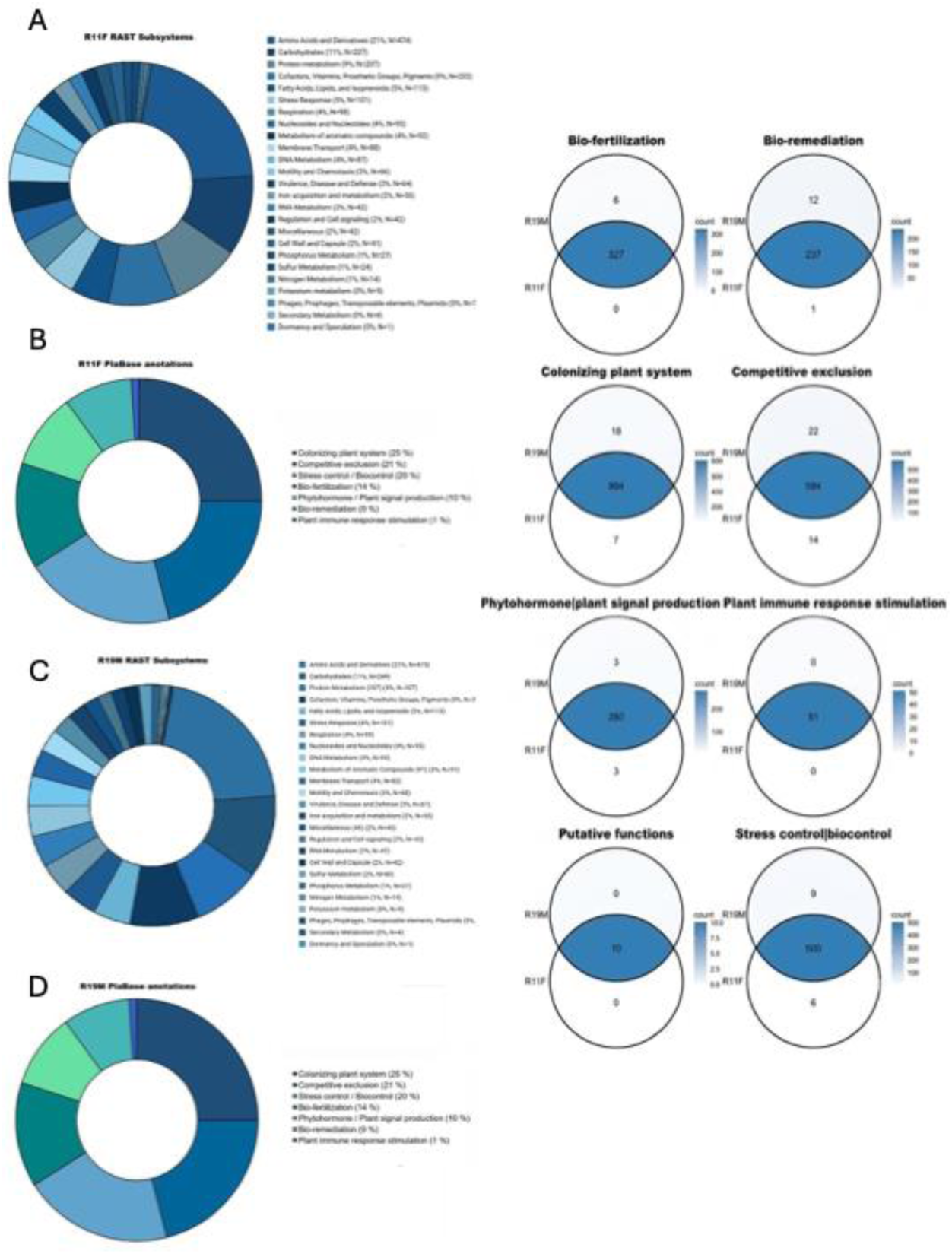
Comparison of the functional annotations of the *Pseudomonas palleroniana* R11F and R19M genomes (Left panels). The upper donut charts show the percentage distribution of coding sequences (CDSs) assigned to metabolic subsystems by RAST for (a) R11F and (b) R19M. The lower charts show the classification of plant growth-promoting trait-related genes (PGPTs) identified using PLaBAse for (c) R11F and (d) R19M. Comparative Venn diagrams showing the distribution of plant growth-promoting and environmental adaptation genes in *Pseudomonas palleroniana* strains R11F and R19M (Right panels).

PLaBAse annotation showed a similar functional distribution, with genes mainly associated with plant colonization (25%), competitive exclusion (21%), stress control/biocontrol (20%), and biofertilization (14%), followed by phytohormone production and plant signalling (10%), bioremediation (9%), and induction of plant immune responses (1%) (Figure 5c,d). Krona analysis further indicated that approximately 68% of the annotated genes were associated with indirect plant growth-promoting mechanisms, including plant colonization, competitive exclusion, stress control, biocontrol, and immune response induction, whereas 32% were related to direct mechanisms, including phytohormone production, bioremediation, and biofertilization.

Comparative analysis revealed a high degree of functional conservation between R11F and R19M (Figure 6). Most genes associated with biofertilization, bioremediation, plant colonization, competitive exclusion, phytohormone production and plant signaling, immune response induction, stress control/biocontrol, and putative functions were shared by both isolates. Notably, genes related to plant immune response induction and putative functions were completely conserved, indicating that the core genomic determinants associated with plant growth promotion are largely shared, with only minor differences in their accessory functional repertoires.

**Figure 6.**
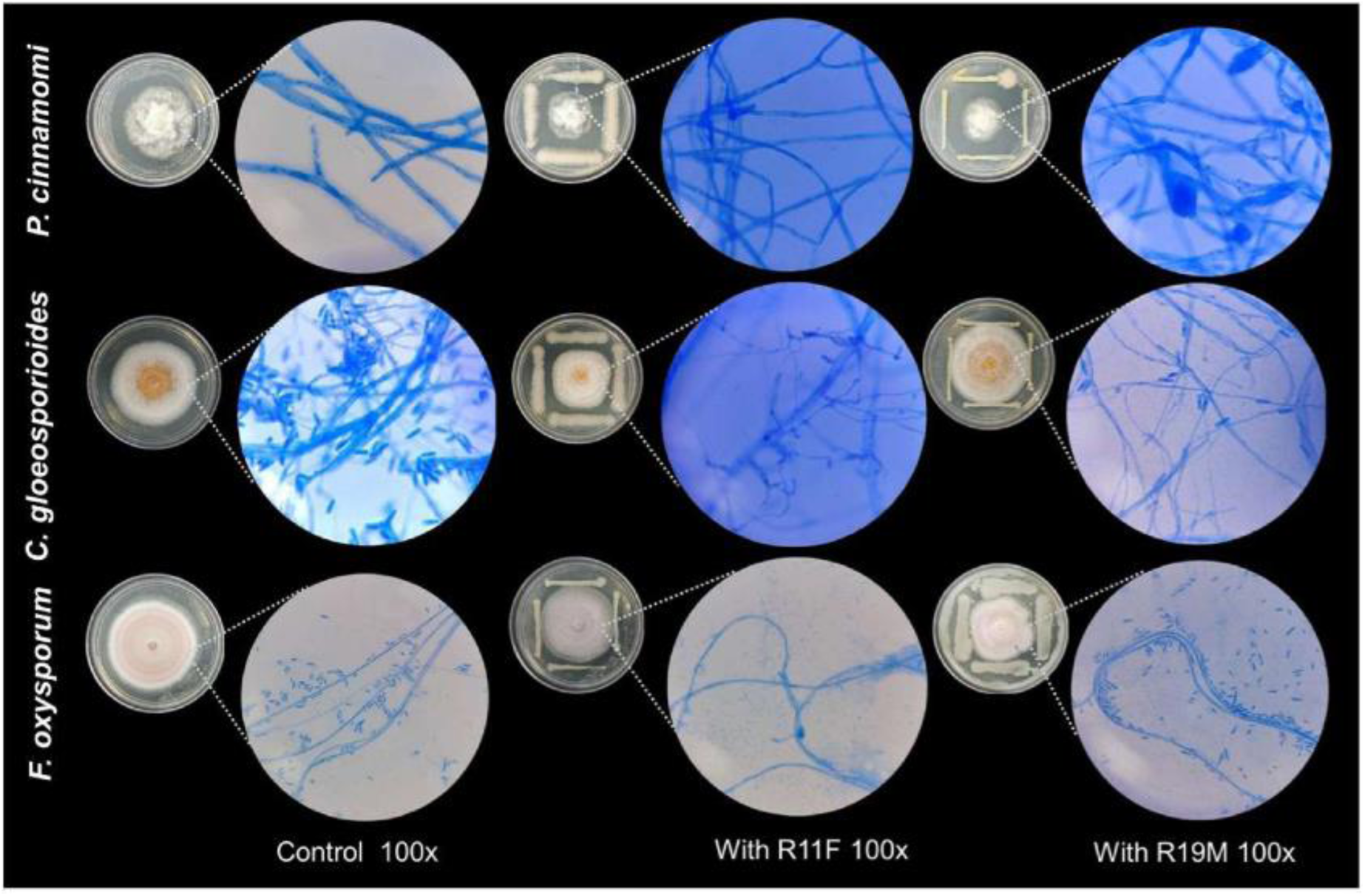
Representative macroscopic and microscopic morphology (100×) of *Fusarium oxysporum, Phytophthora cinnamomi,* and *Colletotrichum gloeosporioides* under control conditions and after exposure to R11F and R19M.

### 3.4 AntiSMASH prediction of secondary metabolites

AntiSMASH analysis predicted 17 and 18 biosynthetic gene clusters (BGCs) in R11F and R19M, respectively (Tables 2 and 3). Both isolates shared BGCs associated with non-ribosomal peptide synthetases (NRPS), siderophores, NRP-metallophores, RiPP-like compounds, arylpolyenes, β-lactones, NAGGN, terpenes, and hydrogen cyanide, suggesting a broad biosynthetic potential for microbial competition, nutrient acquisition, and environmental adaptation. However, strain-specific differences were also identified. R11F harbored an exclusive BGC predicted to produce syringomycin, a non-ribosomal lipopeptide with reported antifungal activity, as well as clusters associated with viscosin and other NRPS metabolites. In contrast, R19M contained NRPS clusters predicted to produce asplenin and kolossin. Overall, these differences indicate partially distinct and potentially complementary biosynthetic repertoires that may contribute to interactions with other microorganisms and the plant environment.

**Table 2.**
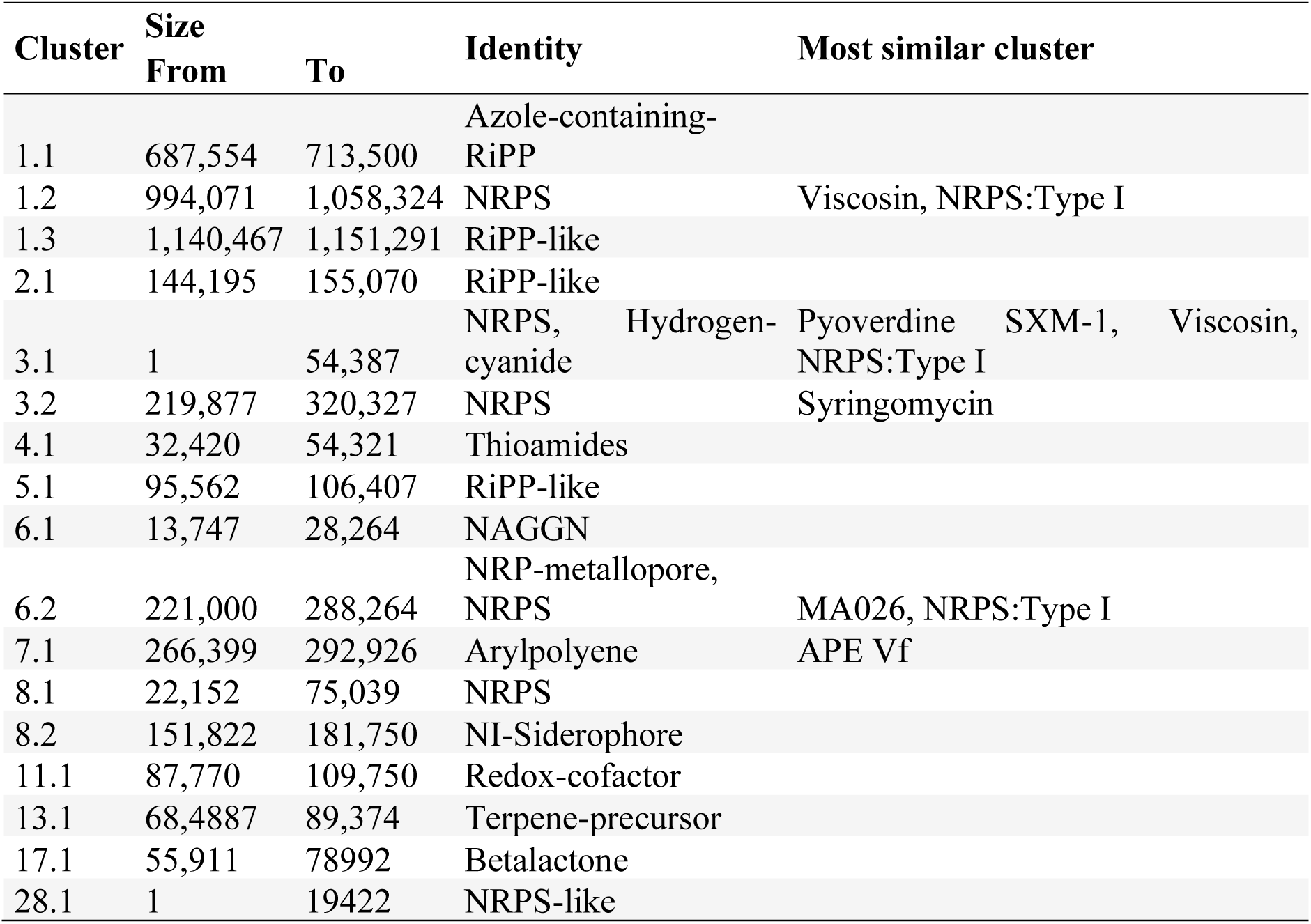
Secondary metabolites predicted by AntiSMASH in R11F.

**Table 3.** Secondary metabolites predicted by AntiSMASH in R19M.

| Cluster | Size<br>From | To | Identity | Most similar |
| --- | --- | --- | --- | --- |
| 16.1 | 107,238 | 118,113 | RiPP-like |  |
| 37.1 | 2,853 | 29,363 | Arylpolyene | APE Vf |
| 46.1 | 1 | 18,698 | NRPS-like |  |
| 58.1 | 818 | 22,965 | Redox-cofactor |  |
| 64.1 | 19,822 | 41,508 | Thiomitides |  |
| 73.1 | 42,221 | 72,149 | NI-Siderophore |  |
| 75.1 | 47,523 | 97,034 | NRPS |  |
| 78.1 | 1 | 14,070 | NAGGN |  |
| 81.1 | 1 | 61,787 | NRP-metallophore, NRPS | Pyoverdine SXM-1, NRPSType I |
| 83.1 | 41,247 | 64,327 | Betalactone |  |
| 85.1 | 10,922 | 21,746 | RiPP-like |  |
| 86.1 | 1,863 | 66,116 | NRPS | Viscosin, NRPS:Type I |
| 89.1 | 45,227 | 71,059 | Azole-containing-RiPP |  |
| 94.1 | 1 | 54,142 | NRPS, Hydrogen-cyanide |  |
| 97.1 | 98,371 | 149,410 | NRPS | Asplenin, NRPS:Type I |
| 98.1 | 1 | 3,091 | NRPS | Kolossin, NRPS:Type I |
| 99.1 | 1 | 45,416 | NRPS |  |
| 102.1 | 68,213 | 89,100 | Terpene-precursor |  |

### 3.5 *In vitro* plant growth-promoting traits

Evaluation of plant growth-promoting traits revealed several functional activities in both bacterial isolates. Nitrogen fixation was indicated by a color change in NFb medium from green to blue, while siderophore production was confirmed by a color change in Chrome Azurol S (CAS) medium from blue to orange, indicating iron-chelating activity. Indole-related compounds were detected using the Salkowski reagent, as indicated by the characteristic reddish colouration. R11F showed the highest production (47.06 ± 1.21 mg/mL), followed by R19M (38.56 ± 7.41 mg/mL), although the difference was not statistically significant (t = 1.1246, df = 2, p = 0.3776). Finally, phosphate solubilization was evidenced by the formation of clear halos around bacterial colonies on Pikovskaya medium, indicating the ability of both isolates to solubilize inorganic phosphate (Supplementary Figure 1).

### 3.1 *In vitro* activity against phytopathogenic fungi

The antagonistic activity of isolates R11F and R19M was evaluated in vitro against three phytopathogens, *Fusarium oxysporum*, *Phytophthora cinnamomi*, and *Colletotrichum gloeosporioides*, over five days of incubation. Both isolates showed a progressive increase in inhibition over time (Table 4, Figure 6). R11F exhibited the highest inhibition of *F. oxysporum* (47.70 ± 1.10%), compared with 40.25 ± 1.11% for R19M. Against *P. cinnamomi*, both isolates showed strong antagonistic activity, with inhibition values of 53.40 ± 0.27% for R11F and 58.42 ± 0.19% for R19M. In contrast, inhibition of *C. gloeosporioides* differed markedly between isolates, reaching 43.78 ± 3.26% for R11F and only 20.76 ± 1.51% for R19M on day 5. Overall, both isolates exhibited antagonistic activity against the tested phytopathogens, although the magnitude of inhibition varied depending on the bacterial isolate–pathogen interaction.

**Table 4.**
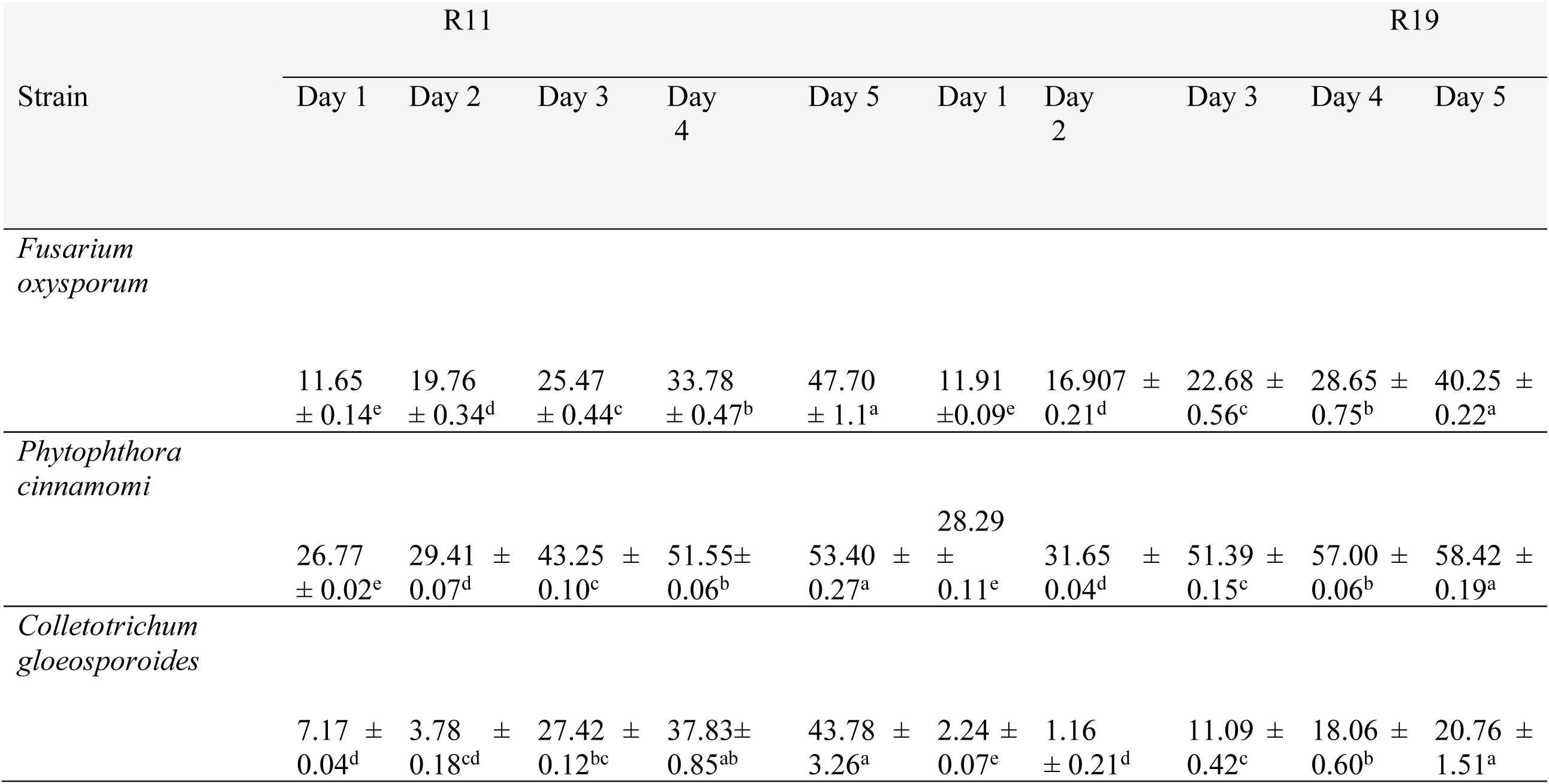
Percentage of mycelial growth inhibition of phytopathogenic fungi by bacterial strains R11 and R19 during a five-day evaluation period.

### 3.2 Effects of bacterial inoculation on plant growth parameters

In tomato plants, R11F significantly increased chlorophyll content and shoot fresh weight compared with the control. However, R19M and the combined R11F–R19M treatment negatively affected root length, shoot length, and the number of lateral roots (Table 5, Figure 7). Significant differences were also observed in root length under R11F and in shoot fresh weight under R19M compared with the control.

**Table 1.** Effects of *Pseudomonas palleroniana* isolates R11F, R19M, and their consortium on the growth and physiological parameters of wheat, lentil, and tomato plants. Values represent the mean ± standard error. Eight replicates were analysed per treatment for wheat and lentil, and five replicates for tomato. Data were subjected to one-way analysis of variance, and means were compared using Duncan’s multiple range test (α = 0.05). Different letters within each column indicate significant differences among treatments. Chlorophyll content in tomato is expressed in arbitrary units. Root and shoot weights in tomato represent fresh weight, whereas those in wheat and lentil represent dry weight. Wheat and lentil were grown in greenhouse pot experiments for 40 days, while tomato plants were grown in a growth chamber for 21 days.

| Phytoparameter | Treatment | Wheat | Lentil | Tomato |
| --- | --- | --- | --- | --- |
| Chlorophyll content<br>$\mu\text{mol/m}^2$ | Control | 475.59 ± 17.90<sup>a</sup> | 292.49 ± 17.96<sup>b</sup> | 10.38 ± 0.74<sup>ab</sup> |
| | R11F | 477.57 ± 16.41<sup>a</sup> | 314.47 ± 15.55<sup>ab</sup> | 12.06 ± 0.54<sup>a</sup> |
| | R19M | 497.87 ± 13.83<sup>a</sup> | 341.58 ± 11.12<sup>a</sup> | 8.34 ± 0.73<sup>b</sup> |
| | R11F – R19M | 513.78 ± 12.29<sup>a</sup> | 339.45 ± 14.93<sup>a</sup> | 9.82 ± 0.79<sup>b</sup> |
| Number of leaves | Control | 15 ± 0.33<sup>a</sup> | 40.25 ± 1.69<sup>a</sup> | 4.44 ± 0.18<sup>a</sup> |
| | R11F | 15.1 ± 0.75<sup>a</sup> | 40.12 ± 2.38<sup>a</sup> | 4.2 ± 0.12<sup>a</sup> |
| | R19M | 15.2 ± 0.41<sup>a</sup> | 41 ± 1.59<sup>a</sup> | 4.36 ± 0.11<sup>a</sup> |
| | R11F – R19M | 15.3 ± 0.61<sup>a</sup> | 41.12 ± 1.89<sup>a</sup> | 4.44 ± 0.17<sup>a</sup> |
| Root length (cm) | Control | 32.54 ± 1.53<sup>b</sup> | 37.86 ± 1.12<sup>a</sup> | 18.99 ± 0.91<sup>ab</sup> |
| | R11F | 38.06 ± 1.51<sup>a</sup> | 30.83 ± 1.38<sup>b</sup> | 19.76 ± 1.09<sup>a</sup> |
| | R19M | 33.08 ± 1.09<sup>b</sup> | 36.2 ± 2.40<sup>ab</sup> | 16.21 ± 1.14<sup>bc</sup> |
| | R11F – R19M | 32.5 ± 1.21<sup>b</sup> | 35.1 ± 2.53<sup>ab</sup> | 14.20 ± 0.98<sup>c</sup> |
| Number of secondary roots | Control | 13.1 ± 0.64<sup>a</sup> | 36.75 ± 2.72<sup>a</sup> | 18.99 ± 0.91<sup>ab</sup> |
|  | R11F | 14.1 ± 0.73<sup>a</sup> | 24.5 ± 1.82<sup>b</sup> | 19.76 ± 1.09<sup>a</sup> |
|  | R19M | 15.4 ± 0.94<sup>a</sup> | 31.63 ± 2.35<sup>a</sup> | 16.21 ± 1.14<sup>bc</sup> |
|  | R11F – R19M | 14.5 ± 0.65<sup>a</sup> | 34.25 ± 1.85<sup>a</sup> | 14.20 ± 0.98<sup>c</sup> |
| Stem lenght (cm) | Control | 43.48 ± 1.20<sup>b</sup> | 33.48 ± 0.60<sup>b</sup> | 18.92 ± 0.91<sup>ab</sup> |
|  | R11F | 46.86 ± 0.69<sup>a</sup> | 34.31 ± 0.96<sup>b</sup> | 19.76 ± 1.09<sup>a</sup> |
|  | R19M | 46.81 ± 0.76<sup>a</sup> | 36.25 ± 1.13<sup>ab</sup> | 16.21 ± 1.14<sup>bc</sup> |
|  | R11F – R19M | 45.87 ± 0.78<sup>b</sup> | 38.7 ± 1.02<sup>a</sup> | 14.20 ± 0.98<sup>c</sup> |
| Shoot dry weigth (g) | Control | 0.89 ± 0.02<sup>a</sup> | 0.47 ± 0.02<sup>a</sup> | 0.06 ± 0.002<sup>b</sup> |
|  | R11F | 0.83 ± 0.04<sup>ab</sup> | 0.46 ± 0.02<sup>a</sup> | 0.12 ± 0.04<sup>a</sup> |
|  | R19M | 0.74 ± 0.02<sup>b</sup> | 0.50 ± 0.02<sup>a</sup> | 0.05 ± 0.003<sup>b</sup> |
|  | R11F – R19M | 0.78 ± 0.04<sup>b</sup> | 0.49 ± 0.02<sup>a</sup> | 0.04 ± 0.003<sup>b</sup> |
| Root dry weigth (g) | Control | 0.66 ± 0.05<sup>a</sup> | 0.66 ± 0.05<sup>a</sup> | 0.03 ± 0.003<sup>a</sup> |
|  | R11F | 0.58 ± 0.03<sup>a</sup> | 0.58 ± 0.03<sup>a</sup> | 0.03 ± 0.004<sup>a</sup> |
|  | R19M | 0.60 ± 0.03<sup>a</sup> | 0.60 ± 0.03<sup>a</sup> | 0.04 ± 0.02<sup>a</sup> |
|  | R11F – R19M | 0.60 ± 0.04<sup>a</sup> | 0.60 ± 0.04<sup>a</sup> | 0.01 ± 0.001<sup>a</sup> |

**Figure 7.**
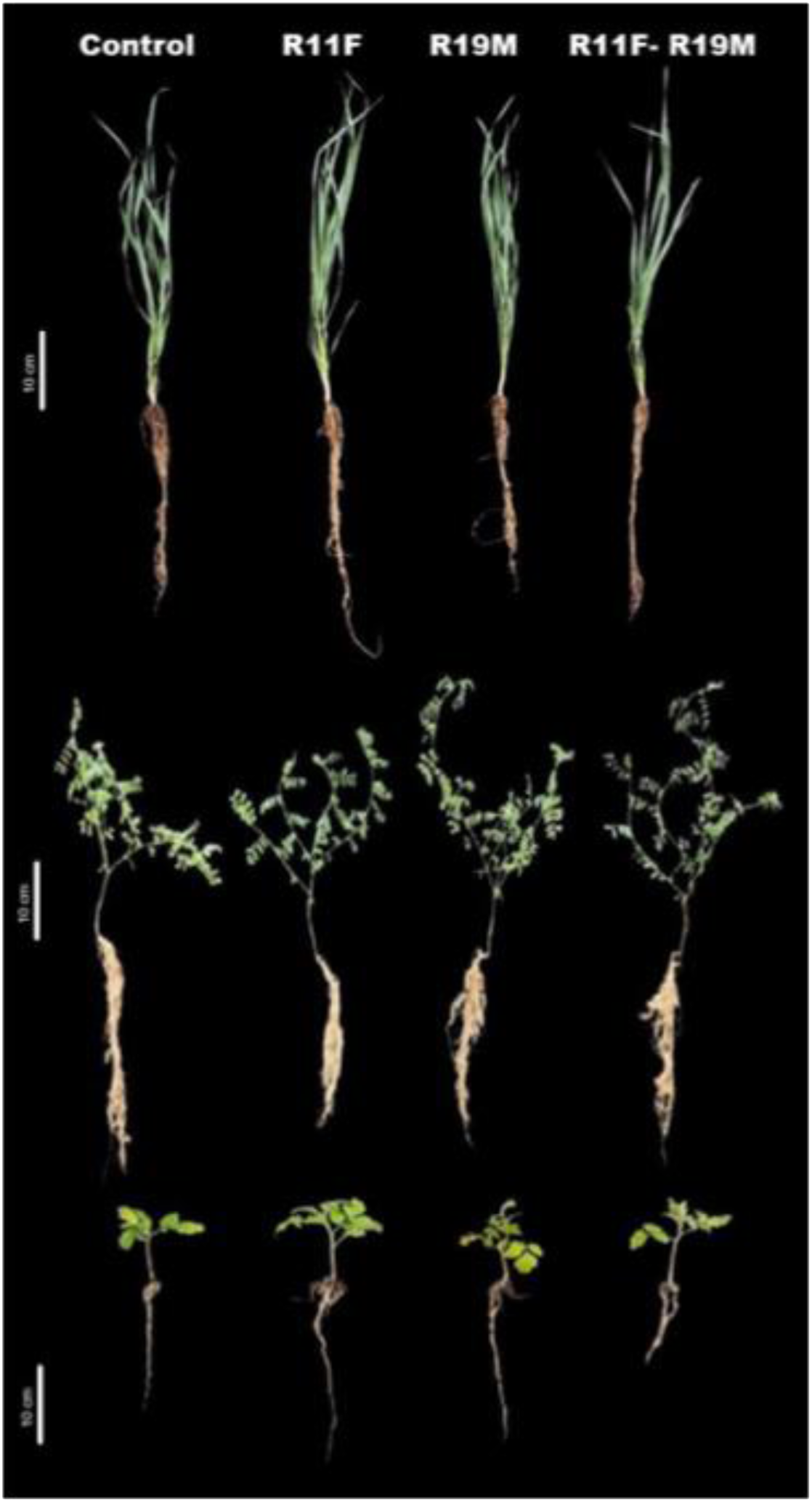
Representative phenotypes of wheat (*T. aestivum*), lentil (*L. culinaris*), and tomato (*S. lycopersicum*) plants grown under control conditions or inoculated with bacterial isolates R11F, R19M, or their consortium (R11F + R19M).

In wheat, R11F significantly increased root length (38.06 ± 1.51 cm) and shoot length (46.86 ± 0.69 cm), whereas chlorophyll content did not differ significantly among treatments. In lentil, bacterial inoculation tended to increase chlorophyll content, with the highest values observed for R19M (341.58 µmol/m²) and the combined R11F–R19M treatment (339.45 µmol/m²). However, R11F significantly reduced root length and the number of lateral roots compared with the control, while shoot length was significantly greater under the combined treatment, reaching 38.7 cm. No significant differences were detected in leaf number or root dry weight in either plant species.

Overall, the effects of bacterial inoculation varied among plant species and treatments, with R11F showing the most consistent plant growth-promoting effects, particularly in tomato and wheat.

## 4. Discussion

The genus *Pseudomonas* has attracted considerable attention because several species exhibit plant growth-promoting and biocontrol activities. For example, *P. fluorescens* LBUM677 and *P. brassicacearum* CDVBN10 have been reported to promote plant growth and increase plant biomass [23–25], whereas *P. protegens* Cab57, *P. chlororaphis*, and *Pseudomonas* sp. P13 exhibit effective biocontrol activity against phytopathogens [26–28]. Other *Pseudomonas* isolates can combine both functions, simultaneously promoting plant growth and suppressing pathogens [29]. Consistent with these reports, R11F and R19M exhibited a genomic repertoire strongly associated with plant–microbe interactions. Approximately 68% of annotated genes were linked to indirect plant growth promotion, including colonization, stress tolerance, immune modulation, and biocontrol, while 32% were associated with direct mechanisms such as biofertilization, bioremediation, phytohormone production, and plant signaling.

The close genomic relatedness of R11F and R19M, together with their isolation from the roots of different crops grown under comparable environmental conditions, suggests that *P. palleroniana* may possess genomic features favoring adaptation to the plant endophytic niche. The conservation of their core genomes may reflect a shared genetic backbone supporting survival and persistence within plant tissues, whereas the strain-specific accessory genes may provide additional functional flexibility to respond to host-specific or microenvironmental conditions. This genomic pattern is particularly relevant for root endophytes, which must tolerate plant-derived stresses while competing successfully within a nutrient-limited and highly selective habitat. The occurrence of closely related but genomically distinct *P. palleroniana* isolates in different plant hosts may therefore indicate a conserved capacity for endophytic colonization coupled with genetic diversification that could facilitate adaptation to particular hosts or ecological conditions. Such genomic plasticity may also contribute to the expression of plant growth-promoting traits and environmental stress tolerance, highlighting the potential of *P. palleroniana* as a versatile plant-associated bacterium [30].

Likewise, previous work demonstrated that *P. palleroniana* Ps006 establishes long-term endophytic colonization more efficiently than *Bacillus amyloliquefaciens*, which was attributed to its repertoire of plant–microorganism interaction genes [31]. Although colonization was not experimentally evaluated in the present study, the enrichment of colonization and competitive-exclusion functions in R11F and R19M supports their potential to establish and persist within plant tissues.

The antagonistic assays further demonstrated that both isolates can inhibit phytopathogens, although the magnitude of inhibition depended on the bacterial isolate–pathogen interaction. R11F showed greater inhibition of *F. oxysporum* and *C. gloeosporioides*, whereas R19M was more effective against *P. cinnamomi* (Figure 6, Table 4). These differences may reflect their partially distinct secondary metabolite repertoires. antiSMASH analysis identified a broad range of biosynthetic gene clusters in both genomes, including those encoding non-ribosomal peptide synthetases (NRPS), siderophores, metallophores, arylpolyenes, RiPP-like compounds, β-lactones, and hydrogen cyanide. NRPS constitute one of the major classes of specialised metabolites produced by *Pseudomonas* [32]. Although both isolates shared several biosynthetic clusters, R11F contained clusters predicted to encode arylpolyene, viscosin, pyoverdine, and syringomycin, whereas R19M contained clusters associated with viscosin, pyoverdine, asplenin, and kolossin. Similar strain-specific variation has been reported among *P. palleroniana* isolates, which may harbour distinct clusters associated with metabolites such as tolaasin, sessilin, putisolvin, pyoverdine, syringomycin, and viscosin [33]. Such differences could contribute to the distinct antagonistic profiles observed in the present study.

Among these metabolites, viscosin may have functions extending beyond direct pathogen inhibition. It has been shown to protect *Pseudomonas* against protozoan predation [34], while Guan et al. (2024) [35] demonstrated that viscosin produced by *P. fluorescens* SBW25 suppressed *Phytophthora* abundance and enhanced root colonization. Thus, the presence of viscosin biosynthetic genes in both R11F and R19M may contribute to both microbial competition and plant colonization. The asplenin-like cluster identified exclusively in R19M may also provide an ecological advantage. Ferrarini et al. (2022) [36] demonstrated that asplenin promotes swarming motility, although its absence did not affect antifungal activity, suggesting a role primarily associated with environmental adaptation.

Pyoverdine was predicted in both genomes and may similarly contribute to bacterial fitness in plant-associated environments. This siderophore is among the most extensively studied metabolites produced by plant-associated *Pseudomonas* species [37]. In addition to facilitating iron acquisition under limiting conditions, inoculation with pyoverdine-producing *P. monsensis* RMC4 has been shown to increase plant biomass under iron deficiency [38]. Pyoverdine has also been associated with enhanced bacterial competitiveness in environments rich in aromatic compounds [39], suggesting an additional ecological advantage for R11F and R19M in the rhizosphere. Both isolates also contained arylpolyene biosynthetic clusters. Although their role in plant–microbe interactions remains poorly understood, arylpolyenes are widely distributed among endophytic and rhizospheric *Pseudomonas* isolates [40] and have been associated with protection against oxidative stress and biofilm formation [41]. The identification of hydrogen cyanide (HCN) biosynthetic genes represents another potential mechanism of antagonism. HCN is a volatile metabolite that has been associated with inhibition rates of up to 57–80% against *Phytophthora infestans* by *Pseudomonas* spp. [42]. Collectively, these genomic features suggest that R11F and R19M possess multiple and potentially complementary mechanisms for microbial competition and pathogen suppression.

Direct plant growth-promoting mechanisms represented a smaller proportion of the predicted functional repertoire. Several *Pseudomonas* species can contribute to biofertilization through siderophore production and phosphate solubilization [23]. According to the PLaBAse annotation, however, R11F and R19M were not particularly enriched in genes associated with phytohormone production or biofertilization, as most of their annotated genes were assigned to indirect plant growth-promoting mechanisms. Previous genomic studies of *P. palleroniana* have reported a broader repertoire of direct plant growth-promoting genes. For example, strain MAB3 contains genes involved in indole-3-acetic acid (IAA) biosynthesis, cytokinin production, and ACC deaminase activity [43]. In the present study, biochemical assays showed that R11F and R19M produced indole-related compounds, solubilized phosphate, and produced a positive reaction in nitrogen-free NFb medium. However, the genomic data did not fully support all of these phenotypes.

No complete IAA biosynthetic pathways were identified in either genomes (Supplementary Table S1), and the Salkowski assay is not specific for IAA. Thus, the indolic compounds produced by R11F and R19M require confirmation using targeted analytical methods. Similarly, although both strains showed a positive response in nitrogen-free NFb medium, the genomes lacked the structural nitrogenase genes (*nifH, nifD*, and *nifK*) and a complete nitrogen-fixation cluster, with only accessory genes such as *nifS* and *nifF* detected. The NFb response may therefore reflect metabolic alkalinisation rather than nitrogen fixation. These findings highlight the importance of integrating phenotypic assays with genomic and targeted molecular analyses to accurately assign plant growth-promoting functions.

In contrast, phosphate solubilization was supported by both phenotypic and genomic evidence. Clear halos were observed on Pikovskaya medium, while both genomes contained a putative glucose dehydrogenase gene (*gcd*) and the *pqqABCDEF* operon, which are involved in gluconic acid-mediated phosphate solubilization (Supplementary Table S1). According to An and Moe (2016) [44], expression of *gcd* and the *pqq* operon constitutes a major phosphate-solubilization mechanism in rhizospheric bacteria such as *P. putida* and is induced in the presence of glucose under phosphate-limiting conditions. Additional genes identified in R11F and R19M, including *phoA*, *phoU*, and *ppx*, may support complementary mechanisms involved in phosphate mobilization through alkaline phosphatase activity and polyphosphate metabolism [45]. Thus, phosphate solubilization appears to be one of the better-supported direct plant growth-promoting traits in these isolates.

Among indirect plant growth-promoting mechanisms, ACC deaminase is particularly relevant because it reduces ACC, the precursor of ethylene, whose excessive accumulation can inhibit plant growth under stress. Both R11F and R19M harbor the *acdS* gene, suggesting a potential role in modulating stress responses in associated plants (Supplementary Table S1). ACC deaminase-producing *Pseudomonas* spp. have been shown to alleviate salinity and drought stress by improving plant physiological performance and biomass [21]. Although ACC deaminase activity was not experimentally assessed here, the presence of *acdS* suggests that R11F and R19M may contribute to plant growth and stress tolerance, particularly under adverse environmental conditions [46]. Similarly, Chen et al. (2024) [47] demonstrated that inoculation with *P. palleroniana* GZNU148 in *Themeda japonica* increased drought tolerance and plant growth while also promoting the recruitment of beneficial rhizosphere bacteria. Whether R11F and R19M can exert similar effects under drought or salinity remains to be determined.

It was demonstrated that inoculation with strain P6 in *Paris polyphylla* increased the production of polyphyllins, metabolites that may be involved in the activation of ISR. However, this potential mechanism remains to be evaluated during the interaction between R11F and R19M, plants, and phytopathogens. Other genomic features may further contribute to stress adaptation. Both isolates contained the trehalose biosynthetic genes *treYZ* and *treS*, which have been associated with osmotic and desiccation stress tolerance in *Pseudomonas.* Orozco-Mosqueda et al. Haga clic o pulse aquí para escribir texto. [48] demonstrated a synergistic effect between trehalose production and ACC deaminase in *Pseudomonas* sp. UW4 inoculated into tomato plants under salt stress. Double mutants deficient in both traits showed reduced survival at 0.2 and 0.8 M NaCl, together with significant reductions in plant growth parameters. Subsequent studies have shown that the trehalose biosynthetic pathways mediated by *treYZ* and *treS* play important roles in osmotic and desiccation stress responses in *Pseudomonas*. Krishna et al. [49] demonstrated that trehalose accumulates under water-deficit conditions, whereas Woodcock et al. [50] showed that mutants deficient in *treYZ* and *treS* exhibited greater sensitivity to salinity and desiccation. Thus, the presence of these genes in R11F and R19M may represent an additional mechanism contributing to bacterial and potentially plant-associated stress tolerance.

In addition to trehalose biosynthesis, R11F and R19M possess other genetic determinants associated with adaptation to osmotic stress. Both genomes contain genes encoding the K⁺ transport systems *kdpABCF, trkAH*, and *kup*, as well as the Na⁺/H⁺ antiporter *nhaA*. These transport systems contribute to ionic homeostasis by facilitating K⁺ uptake and Na⁺ extrusion, thereby supporting bacterial adaptation to saline and osmotic stress conditions. Similarly, induced systemic resistance (ISR) represents another potential mechanism by which *P. palleroniana* may benefit plants.

In addition to trehalose biosynthesis, strains R11F and R19M possess other genetic determinants associated with adaptation to osmotic stress. The genomes of both strains contain genes encoding the K⁺ transport systems *kdpABCF, trkAH*, and *kup*, as well as the Na⁺/H⁺ antiporter *nhaA*. These transport systems contribute to ionic homeostasis by facilitating K⁺ uptake and Na⁺ extrusion, thereby supporting bacterial adaptation to saline and osmotic stress conditions. Wu et al. [22] demonstrated that inoculation with *P. palleroniana* P6 in *Paris polyphylla* increased the production of polyphyllins, metabolites that may be involved in ISR activation. However, whether R11F and R19M can induce similar responses during interactions with plants and phytopathogens remains to be experimentally evaluated.

Beyond plant growth promotion and stress tolerance, the genomes revealed additional features potentially relevant to bioremediation. Both isolates contained a putative *arsRBC* operon, suggesting potential tolerance to arsenic-contaminated environments. ArsR acts as a transcriptional regulator that senses intracellular arsenite and induces operon expression, whereas ArsC reduces arsenate (As⁵⁺) to arsenite (As³⁺), which is subsequently exported through the ArsB transporter. This represents one of the best-characterized arsenic resistance systems in bacteria and has been widely reported in arsenic-resistant microorganisms [51–54].

Both endophytic genomes also harbor *copCD* genes and a CusA/CzcA-family transporter, which are involved in copper uptake and heavy-metal efflux, respectively [55,56]. These features, together with the putative *arsRBC* operon, suggest a potential capacity for heavy-metal tolerance and bioremediation. In addition, plant-associated *Pseudomonas* can contribute to hydrocarbon rhizoremediation through monooxygenases, dioxygenases, and aromatic degradation pathways [57–59]. R11F and R19M harbor genes associated with catechol (*catAC*), protocatechuate (*pcaCDFGH*), and anthranilate (*antABC*) degradation, suggesting the potential to metabolize a broad range of aromatic [60]. Similar pathways have been reported in *Pseudomonas* sp. C11 [61], while *Pseudomonas* inoculation has been shown to simultaneously enhance plant growth and hydrocarbon degradation in plants exposed to n-hexadecane or crude oil [57,62]. These genomic features highlight the potential of R11F and R19M for applications in bioremediation, although their functionality requires experimental validation.

## 5. Conclusion

This study provides integrated taxonomic, genomic, and functional evidence confirming R11F and R19M as *Pseudomonas palleroniana*, with plant growth-promoting traits and genetic repertoire supporting adaptation to abiotic stress. Both isolates showed highly conserved genomes enriched in genes associated with plant colonization, nutrient acquisition, stress adaptation, secondary metabolism, heavy-metal tolerance, and hydrocarbon degradation. Genome mining further revealed diverse biosynthetic gene clusters potentially involved in microbial competition and plant-associated interactions. These genomic features were supported by several functional traits, including siderophore production, phosphate solubilization, indole-related compound production, antagonistic activity against phytopathogens, and plant growth promotion. R11F showed the most consistent plant growth-promoting effects, whereas both isolates displayed pathogen-specific antagonistic activity. Overall, the genomic and functional characteristics of R11F and R19M highlight the ecological versatility of *P. palleroniana* and support their potential as plant-associated bacteria for sustainable agriculture and bioremediation.

## Acknowledgements

G.S. gratefully acknowledge GetGenome (UK) for the high-quality whole-genome sequencing of *Pseudomonas palleroniana* R11F and R19M, which greatly contributed to the genomic analyses presented in this study.

## Figure legends

**Supplementary Figure 1.**
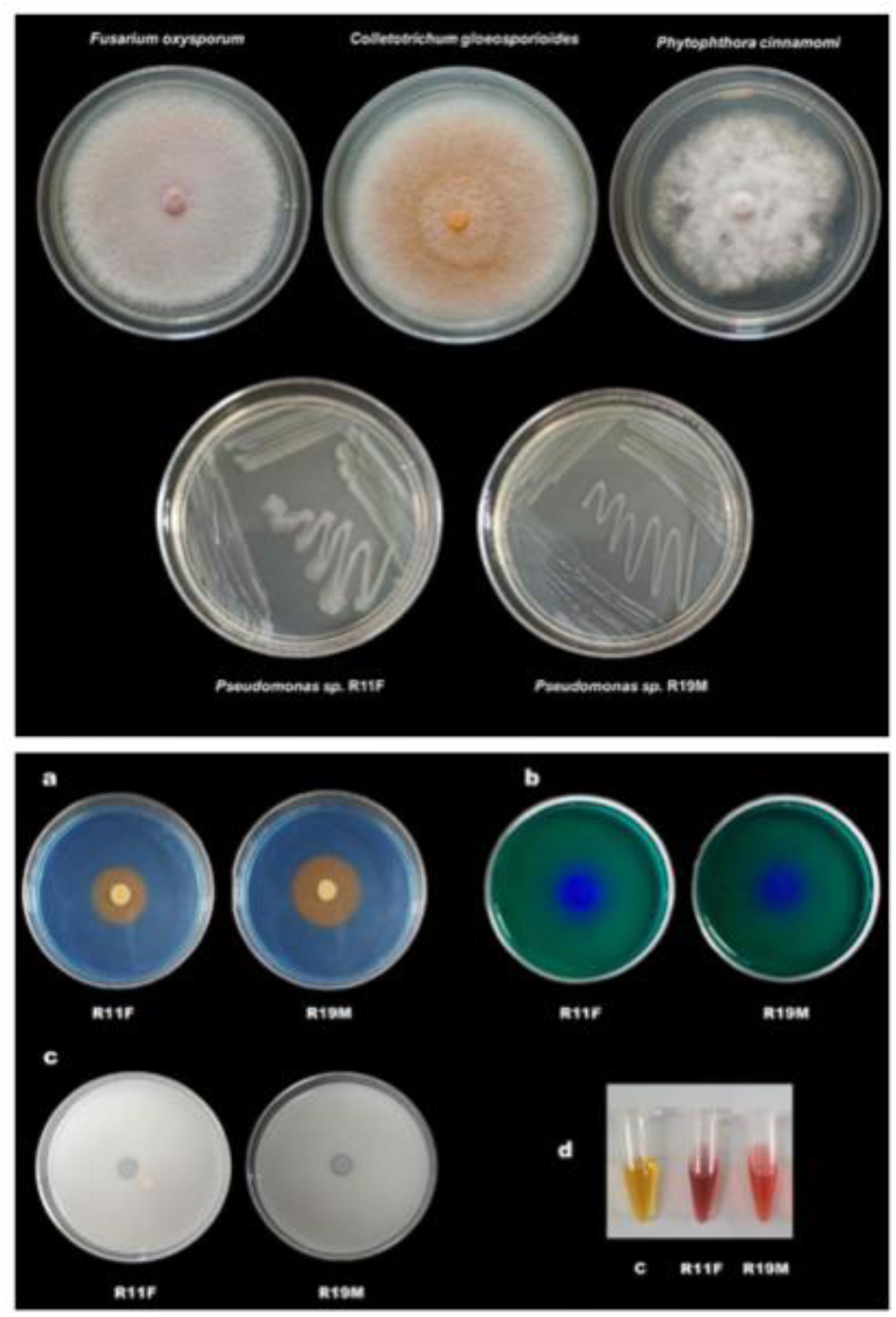
Macroscopic morphology of the microorganisms used in *in vitro* antagonism assays (Upper panel). *In vitro* characterization of plant growth-promoting traits in *Pseudomonas palleroniana* strains R11F and R19M. (Lower panel).

## Notes

### Competing Interest Statement

The authors have declared no competing interest.

