## Supplementary Table 1 for "Unravelling genomic and functional traits of two biocontrol and plant growth-promoting *Pseudomonas* endophytes"

Supplementary Table 1. Shared plant growth-promoting genes identified in the genomes of R11F and R19M.

| **Genes involved in attachment to plant surfaces** | | | | |
|---|---|---|---|---|
| Accession R19M | Accession R11F | Gene | Product | Pathway |
| MGE9873579.1 | MFQ6350311.1 | *iolD* | 3D-(3,5/4)-trihydroxycyclohexane-1,2-dione acylhydrolase (decyclizing) | Myo-inositol degradation |
| MGE9873580.1 | MFQ6350312.1 | *iolB* | 5-deoxy-glucuronate isomerase | |
| MFQ6592184.1 | MFQ6350314.1 | *iolC* | bifunctional 5-dehydro-2-deoxygluconokinase/5-dehydro-2-deoxyphosphogluconate aldolase | |
| MGE9873581.1 | MFQ6350313.1 | *iolE* | myo-inosose-2 dehydratase | |
| **Genes involved in nitrogen fixation and nitrogen metabolism** | | | | |
| Accession R19M | Accession R11F | Gene | Product | Pathway |
| MGE9871269.1 | MFQ6348034.1 | *nifS* | IscS subfamily cysteine desulfurase | Nitrogen fixation |
| MFQ6591915.1 | MFQ6348102.1 | *nifF* | flavodoxin | |
| **Genes involved in phosphate solubilization** | | | | |
| Accession R19M | Accession R11F | Gene | Product | Pathway |
| MFQ6593985.1 | MFQ6347815.1 | *gnl* | 6-phosphogluconolactonase | D-gluconate production |

| | | | | |
|---|---|---|---|---|
| MFQ6590443.1 | MFQ6348403.1 | *ppx* | exopolyphosphatase | Degradation of inorganic polyphosphates |
| MGE9870673.1 | MFQ6348287.1 | *phoB* | phosphate regulon transcriptional regulator PhoB | Phosphate metabolism / Phosphate starvation response |
| MGE9868801.1<br>MGE9870354.1<br>MGE9873764.1 | MGE9868801.1<br>MGE9870354.1<br>MGE9873764.1 | *Putative phoA* | alkaline phosphatase family protein | |
| MGE9871087.1 | MFQ6349742.1 | *pqqD* | pyrroloquinoline quinone biosynthesis peptide chaperone PqqD | PQQ biosynthesis |
| MFQ6590717.1 | MFQ6348733.1<br>MFQ6349745.1 | *pqqA* | pyrroloquinoline quinone precursor peptide PqqA | |
| MFQ6594259.1 | MFQ6349744.1 | *pqqB* | pyrroloquinoline quinone biosynthesis protein PqqB | |
| MFQ6594259.1 | MFQ6349743.1 | *pqqC* | pyrroloquinoline-quinone synthase PqqC | |
| MGE9871086.1 | MFQ6349732.1<br>MFQ6349741.1 | *pqqE* | pyrroloquinoline quinone biosynthesis protein PqqE | |
| MGE9873147.1 | MFQ6349746.1 | *pqqF* | pyrroloquinoline quinone biosynthesis protein PqqF | |
| MGE9873147.1 | MFQ6346281.1 | *Putative gcd* | UDP-glucose dehydrogenase family protein | Gluconic acid |
| MGE9869378.1 | MFQ6346852.1 | *ppc* | phosphoenolpyruvate carboxylase | |
| **Indole synthesis** | | | | |
| Accession R19M | Accession R11F | Gene | Product | Pathway |
| MGE9870282.1 | MFQ6349612.1 | *aspC/ tyrB* | aminotransferase | Indole-3-pyruvate (IPA) pathway |

| | | | | |
|---|---|---|---|---|
| - | - | *ipdC* | Indole-3-pyruvate decarboxylase | |
| MGE9869739.1<br>MGE9870731.1<br>MGE9871785.1<br>MGE9872160.1<br>MGE9872165.1<br>MGE9872169.1<br>MGE9872185.1<br>MGE9872230.1<br>MGE9872299.1 | MFQ6346101.1<br>MFQ6348548.1<br>MFQ6348657.1<br>MFQ6348732.1<br>MFQ6348864.1<br>MFQ6349245.1<br>MFQ6349424.1<br>MFQ6350096.1 | *aldA/aldB* | aldehyde dehydrogenase family protein | |
| - | - | *iaaM* | Tryptophan-2-monooxygenase | Indole-3-acetamide (IAM) pathway |
| MGE9869228.1<br>MGE9869229.1<br>MGE9869527.1<br>MGE9872187.1<br>MGE9872772.1<br>MGE9872975.1 | MFQ6346446.1<br>MFQ6346644.1<br>MFQ6346999.1<br>MFQ6347000.1<br>MFQ6350211.1 | *iaaH* | amidase | |
| - | - | *tdc* | Tryptophan decarboxylase | Tryptamine (TAM) pathway |
| - | - | - | Amine oxidase | |
| MGE9873385.1 | MFQ6346116.1<br>MFQ6348345.1 | *aldD* | aldehyde dehydrogenase | |
| MGE9873864.1 | MFQ6347659.1 | *nit* | nitrilase family protein | Indole-3-acetonitrile (IAN) pathway |
| **Genes involved in response to stress** | | | | |

| Accession R19FM | Accession R11F | Gene | Product | Pathway |
|---|---|---|---|---|
| | MFQ6347641.1 | *acdS* | 1-aminocyclopropane-1-carboxylate deaminase | ACC catabolism |
| | MFQ6350495.1 | *acdS* | 1-aminocyclopropane-1-carboxylate deaminase/D-cysteine desulfhydrase | |
| MGE9868995.1 | MFQ6347227.1 | *kdpA* | potassium-transporting ATPase subunit KdpA | Potassium uptake and osmoregulation |
| MGE9868994.1 | MFQ6347228.1 | *kdpB* | potassium-transporting ATPase subunit KdpB | |
| MGE9868993.1 | MFQ6347229.1 | *kdpC* | potassium-transporting ATPase subunit KdpC | |
| MGE9868996.1 | MFQ6347226.1 | *kdpF* | K(+)-transporting ATPase subunit F | |
| MGE9870556.1 | MFQ6348157.1 | *trkA* | Trk system potassium transporter TrkA | |
| MGE9871468.1 | MFQ6347834.1 | *trkH* | TrkH family potassium uptake protein | |
| MGE9869400.1 | MFQ6346830.1 | *kup* | potassium transporter Kup | |
| MGE9869284.1 | MFQ6346946.1 | *nhaA* | Na+/H+ antiporter NhaA | Ion transport and $Na^+/H^+$ exchange |
| MGE9871188.1 | MFQ6348119.1 | *proB* | glutamate 5-kinase | Biosynthesis and transport of osmoprotectants |
| MGE9870931.1 | MFQ6349907.1 | *proC* | pyrroline-5-carboxylate reductase | |
| MGE9871009.1 | MFQ6349664.1 | *betB* | betaine-aldehyde dehydrogenase | |
| MGE9870319.1 | MFQ6349651.1 | *betT* | choline transporter BetT | |
| MGE9873018.1 | MFQ6346407.1 | *Putative treY* | malto-oligosyltrehalose synthase | Trehalose |

| | | | | |
|---|---|---|---|---|
| MGE9873020.1 | MFQ6346405.1 | *treZ* | malto-oligosyltrehalose trehalohydrolase | |
| **Biorremediation of heavy metals** | | | | |
| Accession R19M | Accession R11F | Gene | Product | Pathway |
| MGE9869234.1<br>MGE9869930.1<br>MGE9870988.1<br>MGE9872465.1<br>MGE9873004.1 | MFQ6346994.1<br>MFQ6348716.1<br>MFQ6348975.1<br>MFQ6349966.1 | *arsR/smtB* | ArsR/SmtB family transcription factor | Arsenic resistance |
| MGE9873078.1 | MFQ6346351.1 | *Putative arsB* | arsenic transporter | |
| MGE9872668.1 | MFQ6346739.1 | *arsC* | arsenate reductase (glutaredoxin) | |
| MGE9872668.1 | MFQ6349159.1<br>MFQ6350468.1 | *copC* | copper homeostasis periplasmic binding protein CopC | Copper resistance |
| MGE9871873.1<br>MGE9872491.1 | MFQ6349158.1<br>MFQ6350467.1 | *copD* | copper homeostasis membrane protein CopD | |
| ACQRBT_05730 | MFQ6349266.1 | *cusA/czcA* | CusA/CzcA family heavy metal efflux RND transporter | Zinc/Cadmium/Cobalt resistance |
| **Aromatic compound degradation** | | | | |
| Accession R19M | Accession R11F | Gene | Product | Pathway |
| MGE9871166.1 | MFQ6351132.1 | *catA* | catechol 1,2-dioxygenase | Catechol degradation |

| | | | | |
|---|---|---|---|---|
| MGE9871165.1 | MFQ6351133.1 | *catC* | muconolactone Delta-isomerase | |
| MGE9869245.1 | MFQ6346985.1 | *pcaC* | 4-carboxymuconolactone decarboxylase | Protocatechuate catabolic genes |
| MGE9869246.1 | MFQ6346984.1 | *pcaD* | 3-oxoadipate enol-lactonase | |
| MGE9869251.1 | MFQ6346979.1 | *pcaF* | 3-oxoadipyl-CoA thiolase | |
| MGE9869249.1 | MFQ6346981.1 | *pcaG* | protocatechuate 3,4-dioxygenase subunit alpha | |
| MGE9869250.1 | MFQ6346980.1 | *pcaH* | protocatechuate 3,4-dioxygenase subunit beta | |
| MGE9869143.1 | MFQ6347085.1 | *pobA* | 4-hydroxybenzoate 3-monooxygenase | |
| MGE9871161.1 | MFQ6351137.1 | *antA* | anthranilate 1,2-dioxygenase large subunit | Anthranilate catabolic genes |
| MGE9871160.1 | MFQ6351138.1 | *antB* | anthranilate 1,2-dioxygenase small subunit | |
| MGE9871159.1 | MFQ6351139.1 | *antC* | anthranilate 1,2-dioxygenase electron transfer component AntC | |

6

7
